# High-throughput measurements of neuronal activity in single human iPSC-derived glutamate neurons

**DOI:** 10.1101/2025.04.07.646449

**Authors:** Isabel Gameiro-Ros, Adam J. Tengolics, Iya Prytkova, Chella Kamarajan, Zhiping P. Pang, Alison M. Goate, Ronald P. Hart, Paul A. Slesinger

## Abstract

Induced pluripotent stem cell (iPSC)-derived neurons provide a promising platform for studying neuronal function and modeling central nervous system (CNS) diseases. However, functional analysis of large populations of iPSC-derived neurons has been challenging. Here, we developed a high throughput strategy targeting N-methyl-D-aspartate receptors (NMDA-R) to enhance neuronal activity and reveal functional phenotypes in human iPSC-induced glutamatergic neurons (iGlut). Using a genetically encoded calcium indicator (GCaMP8f), we first demonstrate that using artificial cerebrospinal fluid (ACSF) lacking Mg^2^+ (Mg^2^+-free) significantly increases neuronal firing, and that firing is enhanced by a potentiator (glycine) but inhibited by the NMDA-R antagonist AP-V. Similarly, multi-electrode array (MEA) recordings also show robust firing in Mg^2^+-free ACSF. Lastly, single-cell patch-clamp electrophysiology confirms the high firing rates in Mg^2^+-free ACSF across multiple iPSC donor lines and also reveals iPSC donor-specific tonic and bursting firing phenotypes. This new methodology provides a scalable, high-throughput method to study neuronal activity in iGlut neurons while preserving single-cell resolution. The strategy also reveals different functional phenotypes, enabling detailed characterization of iGlut neurons in diverse applications such as CNS disease modeling and drug screening. These findings establish a versatile framework for future studies of neuronal network dynamics and individual excitability in iPSC-derived neuronal cultures.

## Introduction

The development of induced pluripotent stem cell (iPSC) technologies in the last two decades has led to unprecedented advances in human CNS disease modeling and drug discovery^1,2^. Human iPSC-derived neurons provide an excellent *in vitro* system to assess neuronal function in physiological and pathological conditions. Human iPSC-based models also overcome many of the limitations of animal models, which cannot replicate species-specific mechanisms^3^ nor incorporate human genetic variants or backgrounds. Directed and transcription factor-based differentiation protocols have been developed to produce relatively pure populations of glutamatergic neurons (iGlut), helping to reduce batch-to-batch variability^4-6^. When co-cultured with glial cells, these iGlut neurons become synaptically mature, exhibiting spontaneous activity^7^. Taking advantage of the iGlut protocols, several groups have used iPSC-derived NGN2-directed glutamatergic neurons to model different human diseases, from neurodevelopmental^8^ and neurodegenerative diseases^9,10^, to other psychiatric diseases such as schizophrenia^11,12^ or alcohol dependence and response^13-15^.

One of the major obstacles in studying the function of iGlut neurons has been their relatively low rate of spontaneous firing^5,7^. In addition, increasing the number of iPSC donor lines to provide sufficient power for statistical comparisons requires the development of high-throughput (HT) assays to monitor neuronal activity. Electrophysiological recordings provide high-content information but are low-throughput, performed on a cell-by-cell basis. Automated patch-clamp technologies are available but require harvesting mature cultures of human neurons in preparation for making recordings, which disrupts neuronal morphology and connectivity. Multielectrode array (MEA) recordings, in which neurons are cultured directly in a well on a plate that has embedded recording electrodes, are another scalable approach, but can be limited by the number of electrodes and location of neurons relative to the electrodes. On the other hand, Ca^2+^ imaging with genetically encoded sensors (e.g. GCaMPs) or Ca^2+^ dyes (e.g. Fluo-4, Fura-2) provides an opportunity to conduct HT measurements of neuronal activity with iPSC-derived neurons. However, the basal activity of neurons, as reflected by Ca^2+^ spikes, is often not substantially different between donor lines, possibly hindered by low spontaneous firing rates. We therefore searched for conditions to improve spontaneous firing. While much focus has been on α-amino-3-hydroxy-5-methyl-4-isoxazolepropionic acid receptors (AMPA-Rs) in iGlut neurons due to their role in mediating excitatory synaptic transmission^16-18^, less is known about the function of N-methyl-D-aspartate receptors (NMDA-Rs) in human neurons. NMDA-Rs are unique because they are inhibited by extracellular Mg^2+^ under physiological conditions and require membrane depolarization to remove Mg^2+^ blockade. If glutamate is bound to the NMDA-R when the neuron is depolarized, NMDA-Rs will support an inward current that excites the neuron.

Here, we investigated the role of NMDA-Rs using Ca^2+^ imaging to assess the activity of a population of human iGlut neurons. We discovered that removing extracellular Mg^2+^ reveals high-frequency neuronal firing, which is highly synchronized in iGlut neurons, and is mediated by NMDA-Rs. We characterized this behavior in multiple iPSC donor lines, replicated the firing behavior with MEA recordings, and confirmed with single-cell whole-cell patch-clamp recordings. Interestingly, we observed unique neuronal firing patterns in Mg^2+^-free ACSF that align with iPSC donor line-specific activity. These results highlight for the first time the usefulness of Ca^2+^ imaging for high-throughput measurements when performed under Mg^2+^-free conditions. This new methodological protocol will enable more efficient population-based measurements of neuronal activity on human iPSC-derived neurons, which can be implemented for drug screening pipelines and phenotypic characterization of iPSC-derived neuronal models of CNS disease.

## Results

### Human iGlut neurons show high frequency and synchronized firing in Mg^2+^-free conditions

To generate a pure population of human iGlut neurons, we used the neurogenin 2 (NGN2) induction protocol with a control donor line (10884) obtained from the Collaborative Study on the Genetics of Alcoholism (COGA) cohort, available through the NIAAA-COGA Sharing Repository^14^. NGN2-derived 10884 neurons were co-cultured with mouse glial cells to promote neuronal maturation and synapse formation^19^. To investigate the activity of these neurons with single-cell resolution while monitoring a relatively large population, we utilized Ca^2+^ imaging in which we expressed lentiviral-transduced GCaMP8f in neurons (human synapsin promoter) at day 28 post differentiation (**Figure 1A**). At approximately 7-10 days post lentiviral infection, green fluorescence could be observed in multiple neurons due to GCaMP8f expression (**Figure 1B**). We measured fluorescent Ca^2+^ spikes in multiple neurons simultaneously using a CCD/CMOS imaging system at 6.1 frames/s with neurons older than 10 weeks post-induction.

**Figure 1:**
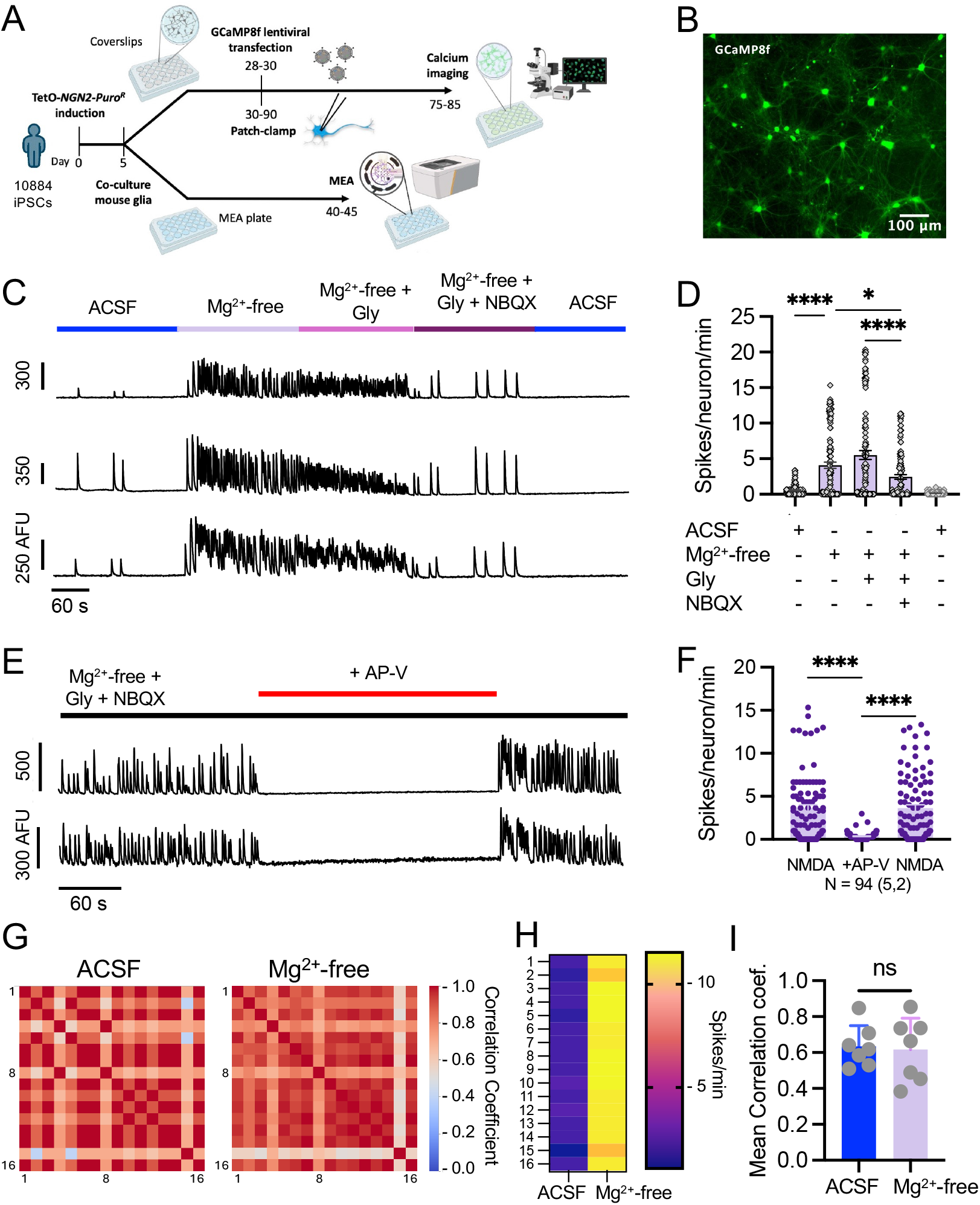
Mg^2+^-free ACSF reveals high firing rates in mature iGlut neurons. **A**) Schematic shows induction and experimental workflow using the NGN2 induction protocol^4,53^, with some modifications (see Methods). GCaMP8f transfection was done on days 28-30 post induction, Ca^2+^ imaging was performed on days 75-85, patch-clamp electrophysiology on days 30-90, and MEA recording on days 40-45 post induction. **B**) Ca^2+^ imaging is used to measure the neuronal activity of the iGlut neurons from line 10884. Representative image shows donor line 10884 iGlut neurons expressing GCaMP8f 2 weeks after lentiviral infection. **C**) ROI fluorescence traces obtained from three different 10884 neurons showing the effects of the indicated buffer conditions. **D**) Plot shows average spikes/neuron/min from two differentiation batches and 115 neurons. Note the significant increase in the frequency of the Ca^2+^ transients in Mg^2+^-free conditions (4.1 ± 0.4 spike/min) compared to ACSF (< 1 spike/min, p < 0.0001, one-way ANOVA, n = 115/2 batches). **E**) ROI fluorescence traces obtained from two different 10884 neurons showing a decrease in Ca^2+^ spikes with NMDA receptor antagonist AP-V (100 µM). **F**) Plot shows average spikes/neuron/min for two conditions (Mg^2+^-free, with 30 µM NBQX and 3 µM Gly; p < 0.0001, one-way ANOVA, n = 94 cells, 2 batches). **G**) High synchrony of neuronal activity observed in both ACSF and Mg^2+^-free ACSF. For the correlation matrices, each cell represents a pairwise correlation between two cells, and the value of the correlation coefficient is represented by the scale bar (Kendall Rank Correlation). **H**) heat map shows firing rate for ACSF and Mg^2+^-free conditions. **I**) Mean correlation coefficient did not change between the ACSF and Mg^2+^-free conditions (n = 7 FOV/2 batches, paired t-test).

To investigate possible NMDA-R-dependent activity, we created an extracellular solution of ACSF that lacks Mg^2+^ (Mg^2+^-free ACSF)^20^. We also created a solution of Mg^2+^-free ACSF that contained the co-agonist glycine (Gly)^21^ and/or a specific ionotropic glutamate receptor antagonist^22^. Under basal conditions with normal ACSF, we observed spontaneously active Ca^2+^ spikes in neurons, but the frequency of Ca^2+^ spikes was relatively low, < 1 spike/min (**Figures 1C, D**). However, exposing the neurons to Mg^2+^-free ACSF led to a significant increase in the frequency of Ca^2+^ spikes (p < 0.0001, one-way ANOVA, n = 115/2 batches), increasing to a mean of 4.1 ± 0.4 spikes/min, with some neurons spiking every 3 s (20 spikes/min). Adding glycine (Gly, 3 µM) to Mg^2+^-free ACSF further increased Ca^2+^ spike activity, reaching an average of 5.5 ± 0.6 spikes/min, with some neurons spiking every 2 s or less. Interestingly, the application of the AMPA-R antagonist NBQX (10 µM) decreased spiking frequency to 2.4 ± 0.3 spikes/min, indicating some contribution of AMPA-Rs to the high firing rates. Ca^2+^ spike activity was reduced further upon the removal of Gly, and the reintroduction of extracellular Mg^2+^ with normal ACSF (**Figure 1C, D**).

The rapid increase in Ca^2+^ spike frequency observed in Mg^2+^-free ACSF with Gly suggested that NMDA-Rs are involved in the high rate of spiking observed. To test this hypothesis, we evaluated neuronal activity in Mg^2+^-free ACSF with an NMDA-R antagonist. We first isolated NMDA-R-dependent activity with Mg^2+^-free ACSF plus glycine (3 µM), and NBQX (30 µM) to selectively block AMPA-Rs (**Figure 1E**). We then tested the effect of the selective NMDA-R antagonist AP-V (100 µM). In the absence of external glutamate, iGlut neurons exhibited a high frequency of spike firing, with an average of 3.6 ± 0.4 spikes/min. The application of AP-V in the Mg^2+^-free ACSF + glycine + NBQX reduced the spike frequency to <0.3 ± 0.04 spikes/min, directly implicating NMDA-Rs in the high-frequency firing. We next performed single-cell RNAseq on iGlut neurons from lines 10884, 8092 and 9206 (**Supplementary Figure S1**). Importantly, we detected mRNA for both AMPA and NMDA receptors (GRID2, GRIA2, GluA2, GRIN2B, GRIA4). We also detected mRNA encoding proteins involved in action potentials and synaptic transmission, including voltage-gated potassium channels (KCND2, KCNQ3, KCNB2), synaptic release proteins (SYN3, RAB3C), voltage-gated sodium channels (SCN2A, SCN3A y SCN9A), voltage-gated calcium channels (CACNA1C) and GABA receptors (GABRB3). These results support the presence of functional AMPA and NMDA receptors, along with the expression of neuronal excitability genes.

In addition to the high rates of spontaneous spiking (**Figures 1C-F**), we observed a high degree of synchronized firing. This can be seen as a large number of spikes that correlate in time in both ACSF and Mg^2+^-free conditions (**Figures 1C, E**). To quantify this, we performed a correlation analysis (Kendall rank correlation) based on the binned (bin size: 1s) time stamps of the Ca^2+^ spikes and plotted the pairwise correlation coefficients (CC) of cells as correlation matrices (10884 is shown in **Figure 1G**). We constructed a correlation matrix for each recording/field of view (FOV) and calculated the mean correlation coefficient (MCC). Both Mg^2+^-free ACSF and ACSF conditions showed high synchrony, regardless of the large difference in firing fate (**Figure 1H**). We averaged the MCCs (**Figure 1I**) and observed a strong positive correlation in both ACSF and Mg^2+^-free conditions (**Figure 1G**). This synchrony did not significantly change in Mg^2+^-free ACSF (n = 7 FOV, paired t-test) (**Figure 1I**). Taken together, these experiments illustrate that iGlut neurons exhibit NMDA-R-dependent activity in external Mg^2+^-free ACSF conditions and are characterized by robust and highly synchronized firing.

### High firing rates of iGlut neurons in Mg^2+^-free solutions are detected with MEA

To see if the increase in neuronal Ca^2+^ spikes measured in Mg^2+^-free ACSF relates directly to changes in electrical activity, we used multi-electrode arrays (MEAs) to electrically measure spiking activity on a faster time scale (MEA). In addition to 10884, we generated iGlut neurons from the iPSC lines of two other donors, 8092 and 9206, to evaluate and compare their behavior under Mg^2+^-free ACSF. For MEA experiments, iGlut neurons were first generated and then co-cultured with astrocytes^7^ on MEA plates (24/48 wells), each equipped with 16 recording electrodes embedded per well. To generate the iGlut neurons we used the same NGN2 protocol, with minor modifications to adapt the procedure to seeding on MEA plates (see Methods). We first recorded basal firing in Mg^2+^-free ACSF for 2 minutes, and then removed the plate, added MgCl_2_ to reach a final concentration of 1.3 mM (i.e., re-create ACSF), and then re-measured the activity in the same plate for another 2 minutes (**Figure 2A**). In this way, we could compare the basal spiking in Mg^2+^-free ACSF with that in ACSF in the same well and recording session. In Mg^2+^-free ACSF, iGlut neurons were highly active in all wells of the MEA plate. After the addition of MgCl_2_, the high basal activity dramatically decreased (**Figure 2B**). The mean firing rate (MFR, **Figure 2C**) was 3.08 ± 0.24 Hz in Mg^2+^-free ACSF and decreased to 0.004 ± 0.001 Hz (n = 48 wells/2 plates, p < 0.0001, Wilcoxon test) after adding MgCl_2_ (normal ACSF). Additionally, inhibiting NMDA-Rs with AP-V in Mg^2+^-free ACSF, completely blocked the activity of the neurons, confirming the involvement of NMDA-Rs in the high firing activity (**Figure 2D**). We compared the weighted mean firing rate (WMFR – mean firing rate base on the electrodes with firing rate greater than 5 spikes/min), number of bursts, and synchrony (described by synchrony index, 0-1) in the three donor lines (10884, 9206 and 8092) with Mg^2+^-free ACSF. The WMFR between the three lines in Mg^2+^-free ACSF did not significantly differ (n_10884_ = 48/two 24 well plates, n_9206_ = 35/one 48 well plate, n_8092_ = 24/one 24 well plate; **Figure 2E**, Kruskal-Wallis test). However, 8092 showed a significantly higher number of bursts compared to 9206; 387.8 ± 49.6 for 8092 and 161.2 ± 20.2 for 9206 (n_10884_ = 48/two 24 well plates, n_9206_ = 48/one 48 well plate, n_8092_ = 24/one 24 well plate; **Figure 2F**, p < 0.001, Kruskal-Wallis test). Donor 10884 showed significantly higher synchrony compared to 9206 (p < 0.0001) or 8092, the synchrony index was 0.74 ± 0.03 for 10884 and ∼0.53 for both 9206 and 8092 (n_10884_ = 48/two 24 well plates, n_9206_ = 39/one 48 well plate, n_8092_ = 24/one 24 well plate; p < 0.05, Kruskal-Wallis test) in Mg^2+^-free ACSF.

**Figure 2:**
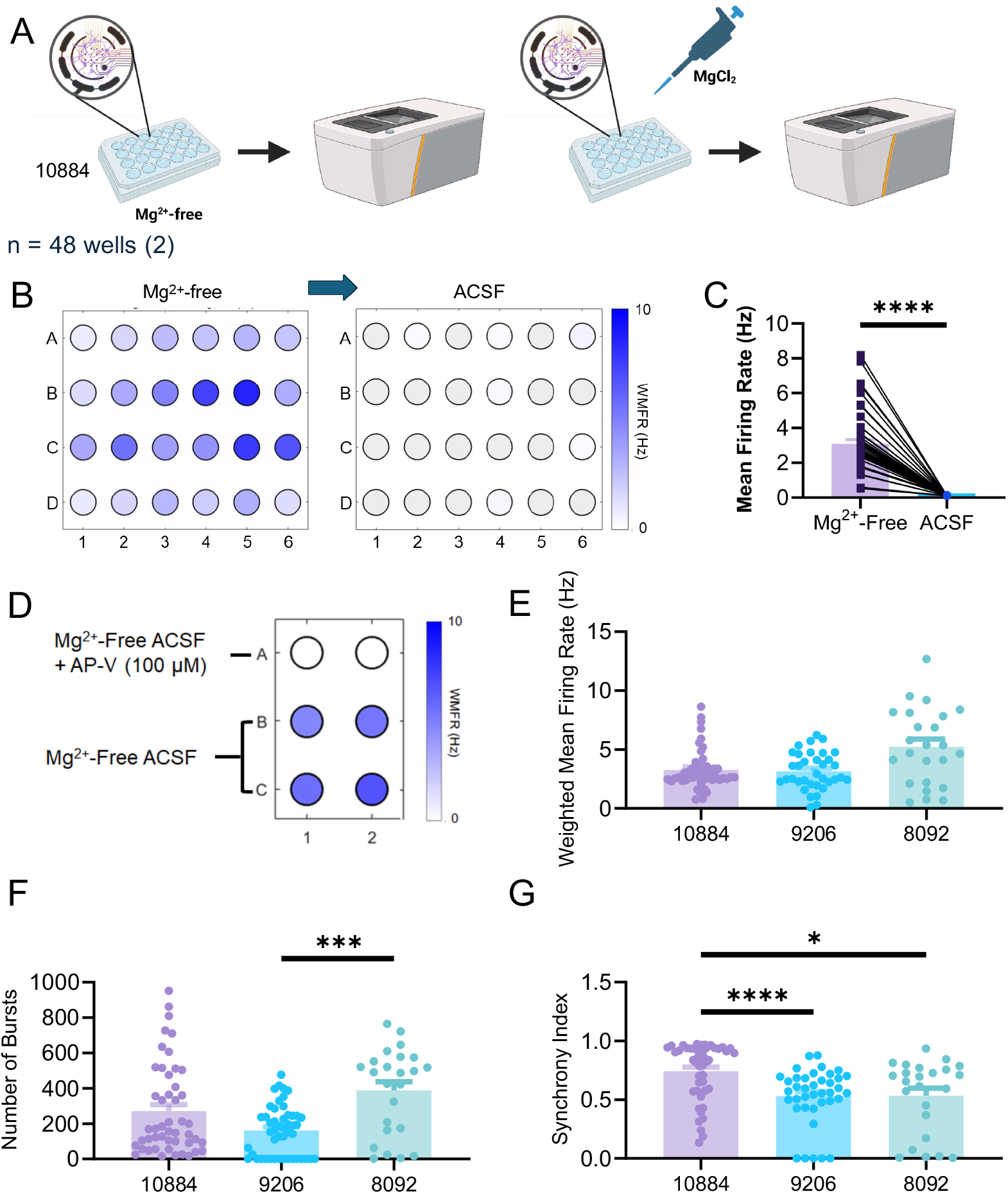
High frequency of firing detected by MEA in Mg^2+^-free condition. **A**) Schematic shows the timeline of the MEA experiment. The activity of iGlut neurons (donor line 10884) was recorded on days 40-45 post induction, first under Mg^2+^-free ACSF, then a MgCl_2_ solution was added to reach a final concentration of 1.3 mM MgCl_2_ and then the activity was measured again in MEA. **B**) Heat map representation of firing activity observed for one 24-well MEA plate before and then after adding MgCl_2_. **C**) Plot shows the MFR changes of donor line 10884 in Mg^2+^-free ACSF (n = 48 wells/2 plates, p < 0.0001, Wilcoxon test). **D)** The heatmap representation shows the WMFR of 6 wells (2 with Mg^2+^-free ACSF + AP-V and 4 wells with Mg^2+^-free ACSF only. **E-G)** Shows three metrics (WMFR, number of bursts and synchrony) and their differences among the three donor lines - 10884, 9206 and 8092 (n_10884_ = 48/two 24 well plates, n_8092_ = 24/one 24 well plate, n_9206_ = 35/one 48 well plate for WMFR, n_9206_ = 48/one 48 well plate for number of bursts and n_9206_ = 39/one 48 well plate for synchrony; *p < 0.05, ***p < 0.001, ****p < 0.0001 Kruskal-Wallis test).

These results demonstrate that the increase in firing rate under Mg^2+^-free ACSF conditions is consistent between two different recording methodologies. Moreover, the high frequency of firing measured with MEAs in Mg^2+^-free ACSF could reveal more robust information from human iGlut neurons derived from different individuals.

### Single-cell electrophysiology confirms high firing behavior in Mg^2+^-free solutions

We next used the gold standard of single-cell whole-cell patch-clamp electrophysiology to further validate the high firing rates of iGlut neurons in Mg^2+^-free ACSF. We generated iGlut neurons from the iPSC lines of three different donors, 10884, 8092 and 9206, using the NGN2-directed differentiation protocol and co-culturing with mouse glial cells. In current-clamp, we first recorded basal firing at the resting potential in ACSF for one minute, and then again in the presence of Mg^2+^-free ACSF. We observed a significant increase in the number of action potentials (APs) in Mg^2+^-free ACSF in iGlut neurons from all three donor lines (**Figure 3A**). The mean AP firing rate increased from ∼5 spikes/min in ACSF to ∼12 spikes/min in Mg^2+^-free ACSF in all three donors iGlut neurons (**Figures 3B, D**). Line 10884 showed an average of < 5 spikes/min in ACSF but > 10 spikes/min in Mg^2+^-free ACSF (**Figure 3D**). Interestingly, while donor lines 10884 and 8092 showed consistent increases in firing with Mg^2+^-free ACSF, 8092 showed both an increase and a decrease in firing rate in different neurons in Mg^2+^-free ACSF (23 % of all recorded cells).

**Figure 3:**
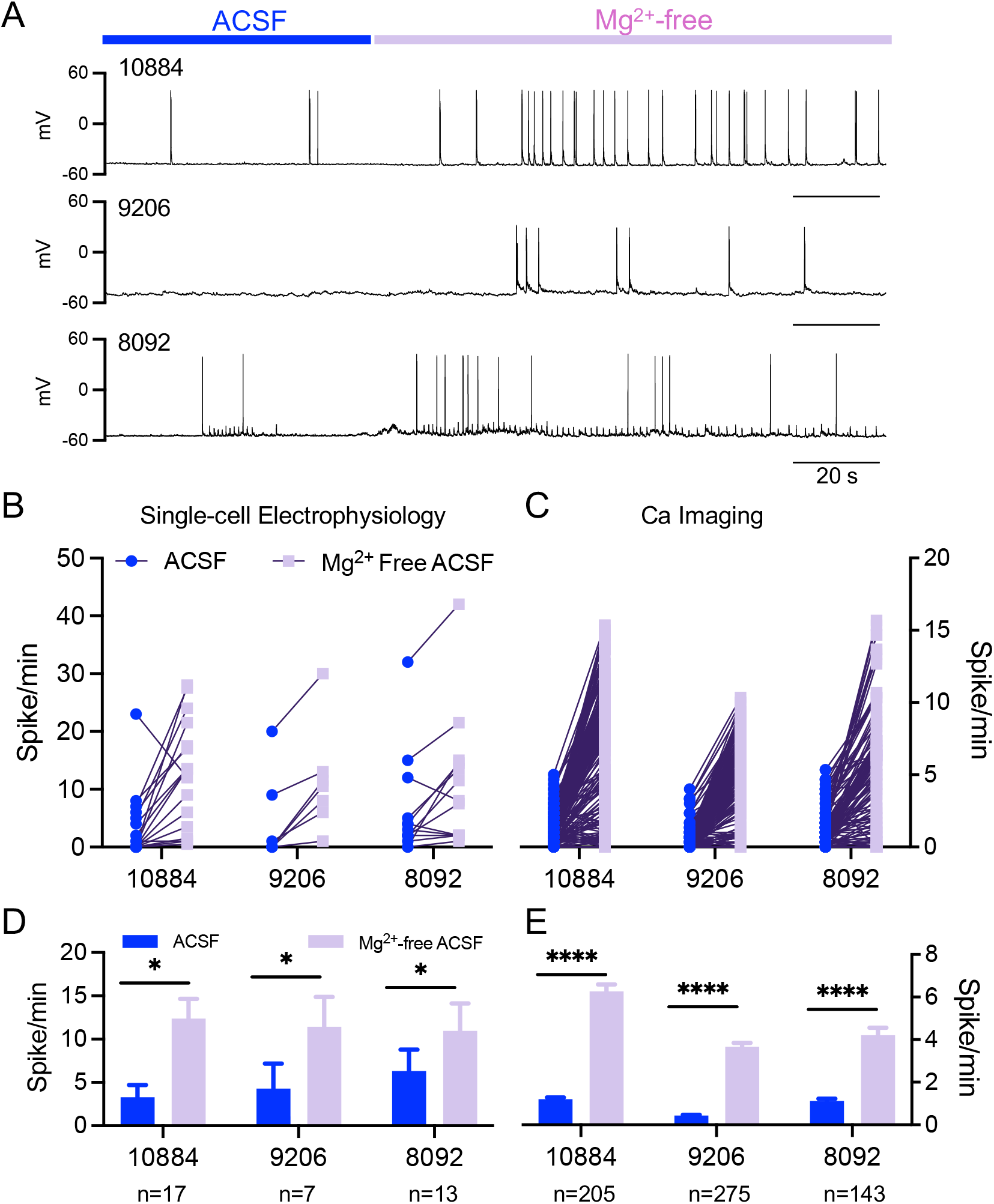
Corroboration of high firing activity in Mg^2+^-free conditions with whole-cell patch-clamp electrophysiology across three iPSC donor lines. **A**) Voltage traces from current-clamp recordings show an increase in firing with Mg^2+^-free ACSF. **B-E**) Graphs show the increase in firing rates (spikes/min) measured by electrophysiology (**B, D**) and Ca^2+^-imaging (**C, E**). Top plots show individual neurons, and bottom plots show the mean ± SEM. There is a significant increase in spikes/min for all three donor lines upon Mg^2+^ removal (10884, 9206 and 8092) (Electrophysiology: n_10884_ = 17/2 batches, n_9206_ = 7/1 batch, n_8092_ = 13/1 batch, p < 0.05, Wilcoxon test; Ca^2+^-imaging: n_10884_ = 205/2 batches, n_9206_ = 275/2 batches, n_8092_ = 143/2 batches, p < 0.0001, Wilcoxon test).

We compared whole-cell patch-clamp recordings with neuronal spiking measured with Ca^2+^ imaging from the same three donor lines. We observed a consistent increase in firing across all three lines in Mg^2+^-free ACSF (**Figures 3D, E**). In Ca^2+^ imaging, the neurons from all three donors showed firing rates ranging from 0.43 ± 0.04 spikes/min for line 9206 to 1.20 ± 0.07 spikes/min for line 10884 (1.1 ± 0.1 spikes/min for line 8092) in ACSF conditions. In Mg^2+^-free ACSF, the calcium spike frequency increased in the three cell lines, showing firing rates of 3.7 ± 0.2, 4.2 ± 0.3, and 6.3 ± 0.3 spikes/min for lines 9206, 8092, and 10884, respectively (**Figures 3D, E**). iGlut neurons from donor line 10884 exhibited the largest response in Mg^2+^-free ACSF in both Ca^2+^-imaging and single-cell electrophysiology. Although there are differences in the temporal resolution between patch-clamp electrophysiology (i.e., ms) and Ca^2+^ imaging (i.e., seconds), there is good concordance between the changes in activity in Mg^2+^-free ACSF in all three lines. The similarity in the results across both modalities also indicates that Ca^2+^ imaging provides a good proxy for measuring spiking frequency in Mg^2+^-free ACSF and is therefore suitable for high-throughput studies of neuronal function.

### Electrophysiological characterization of neuronal activity reveals functional phenotypes

In addition to changes in spiking frequency in Mg^2+^-free ACSF, whole-cell patch-clamp electrophysiology revealed unique firing patterns of iGlut neurons in Mg^2+^-free ACSF. We analyzed the AP firing pattern of individual iGlut neurons from all three iPSC donor lines (10884, 8092 and 9206). We observed that some neurons fired tonically while other neurons exhibited bursts of spikes (two or more consecutive spikes in 100 ms). Interestingly, the distribution of these firing types varied between the donor lines (**Figures 4A-E**). To classify whether a neuron exhibited predominantly bursting or tonic firing activity, we used the distribution of their instantaneous firing rates (IFR) and K-means clustering (see Methods). Each of the three donor lines showed different proportions of neurons that fell into three categories, bursting, tonic, or little or no firing (0-3 spikes) neurons. The distributions of firing types also changed between ACSF and Mg^2+^-free ACSF. In general, there were more low-firing neurons in ACSF for all three donor lines, with an increase in bursting or tonic firing in Mg^2+^-free ACSF. Line 10884 showed the highest proportion of bursting neurons, while line 8092 exhibited the highest proportion of tonic neurons. Line 9206 was overall less active but showed a large increase in tonic firing in Mg^2+^-free ACSF. These differences in firing behavior between lines were not due to differences in intrinsic membrane properties, as there were no significant differences in cell capacitance or resting membrane potential in all three donor lines (p > 0.05, Kruskal-Wallis test, **Figure 4F**).

**Figure 4:**
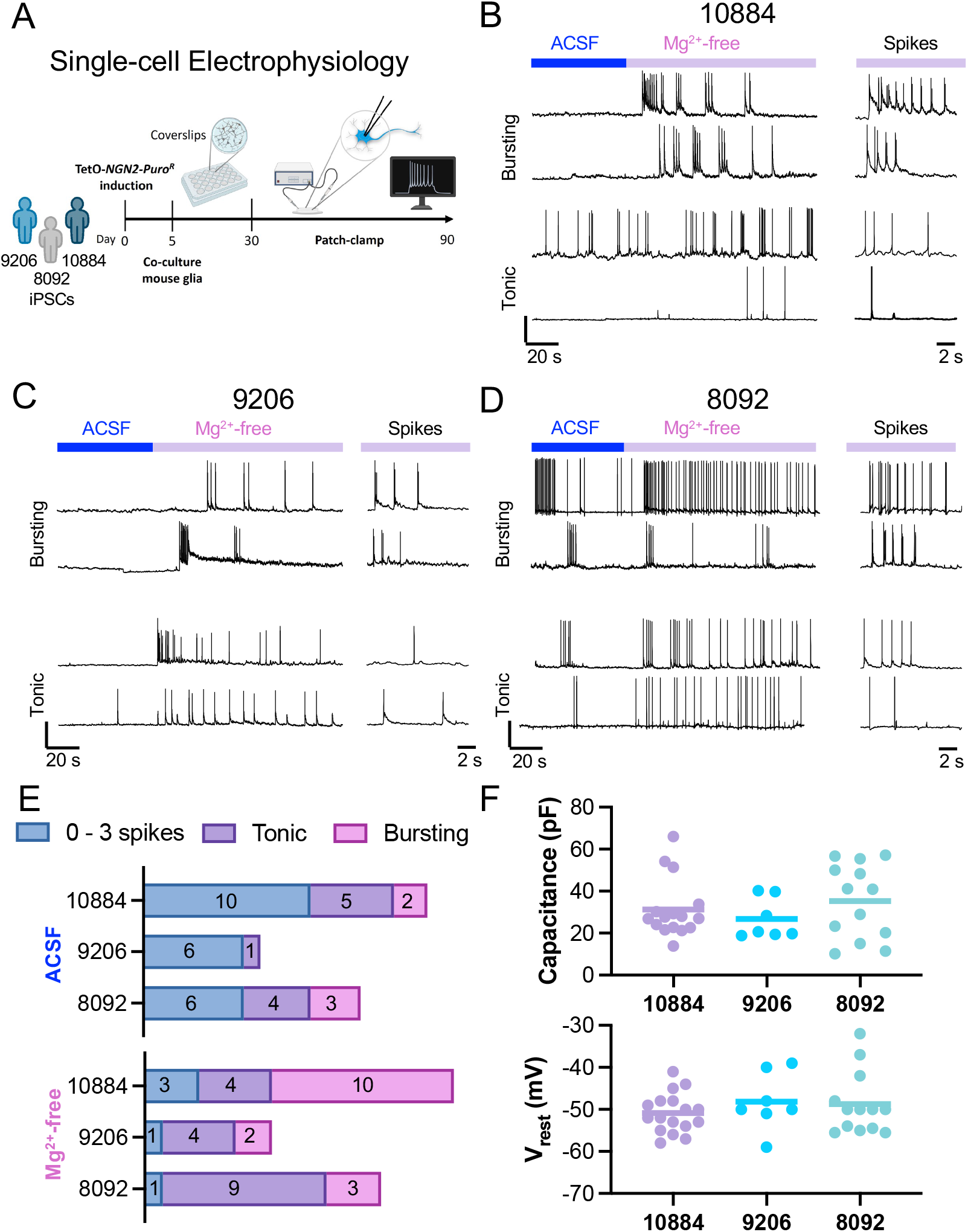
Patch-clamp electrophysiology reveals functional phenotypes for iGlut neurons in Mg^2+^-free ACSF. **A**) Schematic shows patching timeline for all three lines (10884, 9206, 8092). The recordings were carried out on days 30-90 post induction. **B-D**) Left, Representative current-clamp recordings of four different neurons from each donor line showing the effect of switching from ACSF to Mg^2+^-free ACSF. Right – zoom of action potential firing pattern. Scale bars are 40 mV, 20 s and 2 s. **E**) Graphs show the distribution of recordings for little or no activity (0-3 spikes; blue), tonic activity (purple), or bursting activity (pink). Bursts were characterized as 2 or more consecutive spikes in a 100 ms window with a short ISIs. **F**) Graphs show the cell capacitance (pF) and resting potential (mV) for all recordings in the three donor lines. No significant differences were observed (n_10884_ = 17/2 batches, n_9206_ = 7/1 batch, n_8092_ = 13/1 batch, p > 0.05, Kruskal-Wallis test).

These results show that single-cell patch-clamp recordings can provide an additional level of functional characterization when conducted in Mg^2+^-free ACSF, suggesting possible phenotypic differences in iGlut neurons derived from different individuals.

## Discussion

Numerous laboratories and pharmaceutical companies are using iPSC-derived human neurons to study the etiology of various neurological and psychiatric disorders as well as to develop novel therapies^23^. The ability to assess functional differences in human neurons is essential for the successful development of new therapies, as well as addressing the need to scale up studies to look at larger cohorts. Evaluating larger sample sizes in functional studies on iPSC-derived neurons in combination with high-throughput experimental approaches would allow the analysis of multiple iPSC donor lines more efficiently. Lacking in the field has been a strategy to study individual neuronal activity in a high throughput fashion. Here, we describe a new protocol consisting of removing extracellular Mg^2+^ in the ACSF that addresses these limitations in iGlut neurons and supports population-based functional studies of complex neuropsychiatric diseases, such as those having a polygenic contribution^13,24-26^. We show high rates of spontaneous activity in Mg^2+^-free ACSF in three different assays, Ca^2+^ imaging, MEA, and single-cell electrophysiology, and in iGlut neurons from three different donors. Importantly, the high-frequency and synchronized firing observed is NMDA-R-dependent, since it was completely silenced in the presence of the NMDA-R selective antagonist AP-V, and is promoted by the co-agonist glycine. The use of Mg^2+^-free ACSF also uncovered potential unique functional phenotypes for iGlut neurons when assayed with patch-clamp electrophysiology, though this technique is low-throughput as compared to MEA or Ca^2+^ imaging. Furthermore, single-cell RNAseq confirmed the Glu receptor-dependent neuronal activity in all three donor lines under study. Since alcohol (ethanol) blocks the NMDA receptor activity^27^, which contributes to both acute and long-term effects of alcohol as well as the development of alcohol addiction, our findings could have important implications for understanding alcohol addiction and other related disorders.

Automated patch-clamp electrophysiology can overcome the low throughput of classic single-cell patch-clamp electrophysiology. Recent advances in automated patch-clamp have accelerated the study of ion channels and ionotropic receptors expressed heterologously in non-neuronal cells (e.g., HEK-293) for drug screening^28^. However, automated patch-clamp is not readily implemented with mature human neurons. For these chip-based recordings, neurons are harvested and resuspended, which removes synaptic connections, as well as axons or dendrites^29^. Automated patch clamp also requires uniform cell suspensions and purification steps (i.e. cell sorting) to avoid confounding results, which can further disrupt cell structure and viability^29^. Maintaining neuronal morphology and circuitry in iPSC-derived neuronal cultures is essential for an accurate assessment of neuron functionality. Another population-based electrophysiological approach is to use MEAs, which enable the evaluation of neuronal network activity at different time points without disrupting synaptic connections^30^. MEAs have been successfully used to assess iPSC-derived neuronal activity in a variety of diseases^31,32^. However, MEAs do not capture the function of individual neurons in the population. More sophisticated approaches such as the Optopatch platform^33^ combine channel rhodopsin-based actuators to elicit neuronal activity through photonic stimulation with genetically encoded voltage indicators (GEVIs) to monitor action potential firing, leading to a spatially resolved all-optical electrophysiology. Interestingly, the Optopatch technology has been successfully implemented in iPSC-derived neurons^34,35^, including in the study of amyotrophic lateral sclerosis^36^. However, the complexity of this technique makes implementation on a large scale less feasible.

On the other hand, Ca^2+^ imaging provides an excellent approach to studying the neuronal activity of large populations of neurons. Genetically encoded calcium indicators (GECIs), such as GCaMPs, are well-suited for studying neuronal activity at single-cell resolution^37^ and can be selectively expressed in subtypes of iPSC-derived cell types using a cell-specific promoter, offering extended applicability on mixed-cell cultures and organoids. GEVIs), such as QuasArs, and GECIs have been extensively used in the study of neuronal function as a direct measure of action potential firing (GEVIs)^38-40^ or indirectly (GECIs)^41-43^.

### Pros and Cons of HT studies in Mg^2+^-free conditions

Implementation of a protocol that compares ACSF to Mg^2+^-free ACSF affords several advantages over previous methods. First, Ca^2+^ imaging can now be used to monitor hundreds of neurons simultaneously, including in mixed cultures with other cell types. Calcium imaging can assess the activity of iPSC-derived iGlut neurons with single-cell resolution within a population, while maintaining crucial neuronal connections. Second, MEA measurements in Mg^2+^-free ACSF align well with those obtained using Ca^2+^ imaging. Although it does not offer single-cell resolution, recording neuronal activity in Mg^2+^-free ACSF is straightforward for MEA and allows for pair-wise comparisons of Mg^2+^-free ACSF vs ACSF conditions. Third, in addition to GECIs like GCaMP, chemical Ca^2+^ dyes (e.g. Fluo-4) can also be used to monitor Ca^2+^ spikes in Mg^2+^-free ACSF. However, the ability to specifically target neurons with chemical dyes is not possible, since these indicators are not cell type specific like their GECI counterparts. Lastly, although it provides slower throughput, single-cell patch-clamp electrophysiology reveals a rich panoply of firing behaviors for iGlut neurons across different neurons, including intrinsic and subthreshold membrane potential changes as well as synaptic activities. Some disadvantages of this Ca^2+^ imaging approach are that large imaging datasets are generated, which require automated spike detection software. Another limitation of this approach is that the Mg^2+^-free ACSF protocol described might only apply to iGlut neurons, where the high-frequency and synchronized firing observed is NMDA-R-dependent. Lastly, the temporal resolution of Ca imaging is slower than patch-clamp electrophysiology, although we show good agreement between the two methods in Mg^2+^-free ACSF.

In summary, this study establishes a high-throughput methodology to enhance the functional characterization of human iPSC-derived neurons under Mg^2^+-free conditions. By leveraging Ca^2^+ imaging, we demonstrate that Mg^2^+-free ACSF robustly increases spontaneous neuronal firing in iGlut neurons, is reversible, and is population analyses of neuronal activity across multiple iPSC donor lines. This approach not only overcomes challenges associated with low spontaneous activity in iPSC-derived neurons but also reveals donor-specific functional phenotypes, highlighting its utility in personalized CNS disease modeling and drug screening. These findings provide a foundation for integrating high-throughput functional studies into large-scale iPSC-based research efforts.

## Materials and methods

### iPSC culture and NGN2 induction

We used a neurogenin2 (NGN2) protocol to generate glutamatergic neurons (iGlut)^4^, with some modifications. iPSCs were selected from three different COGA study donors who were unaffected by AUD: lines 10884 (male, 27 y/o), 8092 (female, 33 y/o), and 9206 (female, 22 y/o) (see the l COGA repository for full details and characterization data upon request^44^). Each donor line was tested for expression of pluripotency markers and for the presence of a euploid karyotype^13^. Original donor iPSC lines received from the COGA repository had the following passages: P7 for line 9206, P14 for 8092, and P11 for 10884; they were banked with 2-5 additional passages, and used for the experiments described in this manuscript with 4-6 additional passages from the original vial. The human iPSCs donor lines were banked using culture media (StemMACS™ iPS Brew XF, human, Miltenyi Biotec, cat. #130-104-368) with 10% DMSO (Sigma-Aldrich, cat.# D8418; 0.5-1*10^6^ cells/cryotube) and frozen in liquid nitrogen, and were thawed and cultured on Matrigel-coated (1:250 dilution; Corning Inc. cat. #354230) 6-well plates for at least one week before the induction. All donor iPSC lines in culture were checked daily for bacterial and/or fungal contamination (bright field microscope visualization) and tested once a month for mycoplasma contamination. iPSCs donor lines were maintained in StemMACS™ media until the day of induction and passaged at least once before starting the differentiation, with a routine splitting rate of 1:6, using ReLeSR™ as dissociation reagent (Stemcell Technologies, cat. #100-0483).

For generating the iGluts we used a doxycycline-inducible tetO-Ngn2-T2A-Puro/rtTA lentiviral induction. Lentiviral plasmids to induce neurons (pTet-O-Ngn2-puro and FUdeltaGW-rtTA) were obtained from AddGene (#52047 and #19780, respectively), sequenced to confirm identity, and packaged using standard packaging plasmids (pMD2.G, #12259; pMDLg/pRRE, #12251; pRSV-Rev, #12253). The hSyn1-driven GCaMP8f virus was constructed from pGP-AAV-syn-jGCaMP8f-WPRE (AddGene #162376) by cloning the hSyn1-gGCaMP8f cassette into a backbone made from FUGW (#14883). The resulting plasmid is available from AddGene with cat. #197034. On the initial day of induction (D0) we dissociated the iPSCs with Accutase™ (Stemcell Technologies, cat.# 07920) and re-plated them in StemMACS™, supplemented with the NGN2 and rtTA containing lentiviruses and 5 µM of the ROCK inhibitor Y27632 (RO kinase inhibitor, Miltenyi Biotec cat.# 130-103-922), onto another Matrigel-coated (2:250 dilution) plate, in a specific cell density (2.5-3*10^5^/ml). The cells were counted with a Countess 3 automated cell counter (Invitrogen). On day 1 (D1), we changed the media to Neurobasal™ (Thermo Fisher/Gibco, cat. # 21103049), supplemented with 1 V/V% CultureOne™ (Thermo Fisher/Gibco, cat. # A3320201), 2 V/V% B27™ (Thermo Fisher/Gibco, cat. # 17504044), 1 V/V% GlutaMAX™ (Thermo Fisher/Gibco, cat. #35050061), and 0.1 V/V% ascorbic acid, with 2 µg/ml Doxycycline to activate the Tet-ON-based NGN2 induction. On D2, we added 1 µg/ml of Puromycin in the culture media for 48 h to select for transduced cells. On D4, we removed the Puromycin but maintained Doxycycline for another 24 h. In parallel, on D4 we plated the mouse glia on acid-edged, Matrigel-coated (4:250 dilution) coverslips (12 mm) in a 24-well plate. Neurobasal was supplemented with 5 V/V% Heat Inactivated Fetal Bovine Serum (HI FBS, Fisher Scientific, cat. # MT35011CV) as a plating media for the mouse glial cells (6.5-7*10^4^ cells/well). On D5, iGlut neurons were re-plated onto the mouse glia in the 24-well plate. First, iGlut were dissociated with Accutase (6 minutes incubation at 37°C), harvested and diluted in DMEM (Thermo Fisher, cat. #11965092, 1:4 dilution), then centrifuged for 4 minutes at 0.8 rcf at room temperature. The pellet was resuspended in 1 ml of Neurobasal + 2 V/V% FBS, counted and neurons diluted to 1-1.5*10^5^ cells/ml in a 24 ml final volume of Neurobasal + 2 V/V% FBS. We gently aspirated the media from mouse glia and plated the iGlut neurons (1 ml of neuron suspension to each well) onto the glia. On D8, we switched the coculture to Neurobasal plus™ media (Thermo Fisher/Gibco, cat. #A3582901) supplemented with 1 V/V% CultureOne™ (Thermo Fisher/Gibco, cat. #A3320201), 2 V/V% B27™ Plus (Thermo Fisher/Gibco, cat. #A3582801), 1 V/V% GlutaMAX™ (Thermo Fisher/Gibco, cat. #35050061), 0.1 V/V% ascorbic acid, 2 V/V% FBS and 2 µM Ara-C (to prevent survival of mitotic cells) by doing a half media change with Neurobasal plus + 4 µM Ara-C. On D11, we refreshed the Ara-C with another half media change with 2 µM Ara-C in Neurobasal plus + 2 V/V% FBS. On D15, we started the removal of Ara-C (half media change only with Neurobasal plus + 2 V/V% FBS) and then cultures were maintained with half media changes using Neurobasal plus + 2 V/V% FBS twice a week.

### Mouse glia cultures

We used postnatal day 3 C57BL/6J wild-type mice to produce glia cell cultures. Mice were housed with their breeding pairs of two females and one male in a 12 h light/dark cycle at 22 ± 2°C with food and water available *ad libitum*. All mouse procedures were carried out following protocols approved by the Institutional Animal Care and Use Committee at the Icahn School of Medicine at Mount Sinai. Dissected brain cortices from 3 pups (at p0-3) were dissociated in a papain-containing (Sigma-Aldrich, cat. #A3582801) solution (19-38 units Papain, 0.5 μM EDTA, 1 μM CaCl_2_ in HBSS) at 37°C for 15 minutes, with gentle shaking every 5 minutes. The dissociation solution was removed and treated tissue was washed twice (the media was added then removed with caution) with 10 ml MEF media (88 V/V% DMEM, 10 V/V% calf serum (Fisher Scientific, cat. #SH3008704), 1 V/V% sodium pyruvate (100 mM, Thermo Fisher, cat. #11360070), 1 V/V% 100x MEM non-essential amino acid solution (Thermo Fisher, cat. #11140050), 0.0008 V/V% 2-mercaptoethanol (Sigma-Aldrich, cat. #M6250)). The tissue was then triturated in 1ml of MEF with a pipette until no large tissue chunks were visible. An additional 4 ml MEF media was added, and the mixture was passed through a 0.4 μm cell strainer into a 50 ml falcon tube containing 5 ml MEF media and seeded in T75 flask. The media was changed the next day and then changed again every 3 days until the glial cells became confluent (in approximately 7 days). The cells were passaged at least once using trypsin and MEF media, with the media being changed every 3-4 days until plating on coverslips for the neuronal co-cultures. We’ve never used glia older than 10 days or passaged more than three times (more than P3).

### scRNAsequencing

Single-cell RNA sequencing was performed on induced neuron cell villages as described previously^14,46^. Cells were matched with subjects using demuxlet^47^ and aggregated by subject. Scaled, normalized gene counts were extracted and plotted using the pheatmap function in R^48^ to plot color-scaled expression of genes shown from donor lines 10884, 9206, and 8092; genes were grouped by function. Raw single cell RNAsequencing data is publicly available on the GEO repository, with accession number GSE293298.

### Ca^2+^ imaging

Lentivirus expressing GCaMP8f was produced and used to transduce (1*10^6^ IU/100,000 cells) iGlut neurons at D28-D33, achieving expression levels suitable for Ca^2+^ imaging around 2 weeks after infection (D42-D47). We measured Ca^2+^ spikes at D72-D85 (see Figure 1A) using a Nikon Eclipse TE2000-U microscope equipped with 20x objective, a 480 nm LED (Mic-LED-480A, Prizmatix Ltd.) passing through a HQ480/40x nm excitation filter (Q505LP dichroic mirror) and a HQ535/50m emission filter (Semrock), a sCMOS Zyla 5.5 camera (Oxford Instruments, Andor), and NIS elements AR software (version 5.21.03, Nikon) for data collection and analysis. The images were obtained using 150 ms exposure time, 6.14 fps frame rate, and 4×4 binning, under constant perfusion with different ACSF-based solutions, in a laminar flow diamond-shaped chamber (Model #RC-25; Warner Instruments) at room temperature (RT, ∼20°C). For all of our experiments, we used artificial cerebrospinal fluid (ACSF; 125 mM NaCl, 5 mM KCl, 10 mM D-Glucose, 10 mM HEPES-Na, 3.1 CaCl_2_, 1.3 mM MgCl_2_) and Mg^2+^-free ACSF (125 mM NaCl, 5 mM KCl, 10 mM D-Glucose, 10 mM HEPES-Na, 3.1 CaCl_2_) as base solutions. To record only NMDA-R only activity, we used Mg^2+^ free ACSF with 10 µM NBQX (AMPA-R antagonist; Figures 1C, D) or 30 µM NBQX (Figures 1E, F). In some experiments, 100 µM AP-V (NMDA-R antagonist, Figures 1E, F) was also applied to block NMDA-R-dependent activity. We also examined the effect of the NMDA-R co-agonist glycine (3 µM Gly).

For Ca^2+^ imaging data analysis, we selected ROIs (placed on the soma of neurons) and exported raw fluorescence traces. We used a baseline-drift correction with penalized least-squares algorithm (AirPLS)^49^ and calculated ΔF/F_0_ with the following formula [F(t)-F_0_)/F_0_], where F_0_ was the minimum fluorescence intensity (RFU) in the first 10 s of the recording. Prior to peak detection, we applied a 3rd order Butterworth filter using the default “signal” package in R (version 4.3.0)^50^. Ca^2+^ transients were detected by a custom-made R script. Ca^2+^ transients were identified based on their kinetics: < 6.4 s width, more than 325 ms rise phase, more than 650 ms fall phase, the rise phase duration less than the fall phase duration, the peak height is more than 5*SD and more than 5*max background signal^14^.

To measure the synchrony, Kendall rank correlation was performed on the binned (bin size: 1s) timestamps of the Ca^2+^ spikes. We only used cells/ROIs for this analysis, which had at least 3 spikes in both ACSF and Mg^2+^-free conditions. We counted only the significant correlations (p < 0.05) toward the MCC.

### Electrophysiology

We carried out whole-cell patch-clamp electrophysiology with iGlut neurons as described previously^15^. Briefly, we recorded spikes in I=0 current-clamp mode from iGlut neurons derived from three representative human donor lines, 10884, 9206, and 8092. Borosilicate glass capillary pipets (3” thinwall, 1.5 OD/1.12 ID, World Precision Instruments) were pulled with a Narishige PC-10 puller (Narishige International USA) and had resistances of ∼4 MΩs, with K-D-Gluconate internal solution (140 mM K-D-Gluconate, 4 mM NaCl, 2 mM MgCl_2_-6H_2_O, 1.1 mM EGTA, 5 mM K-HEPES, 2 mM Na_2_ATP, 5 mM Na-Creatine-PO_4_, 0.6 mM Na_3_GTP) and ACSF external. Recordings were made with a MultiClamp 700B amplifier (Molecular Devices), low-passed filtered at 2 kHz, digitized at 20 kHz with a Digidata 1440A A/D converter (Molecular Devices) and stored on a laboratory computer. Recordings were carried out with pClamp 10 and analyzed with Easy Electrophysiology software. Before the AP counting, we removed the baseline based on the first 30 s of the recording and a polynomial fitting method to remove the possible baseline fluctuation, which was crucial for the AP thresholding. For spike counting, we used automatic thresholding, which uses the first derivative method, with the rise time 3mV/ms, fall time 1mV/ms, and 5 ms AP width.

We classified iGlut neurons as *tonic* and *bursting* neurons based on the interspike intervals (ISIs). First, we determined the instantaneous firing rate (IFR = 1/ISI) and then calculated the standard deviation of the IFRs. Cells with tonic firing usually present single APs with relatively equal ISIs between them, while bursting neurons have short ISIs between the spikes in the bursts and long ISIs between the bursts themselves. Based on this, we were expecting a high SD of IFR for bursting and a low SD of IFR for tonic neurons. To classify the recorded population of iGlut neurons, we first plotted the IFR SD values for each cell, then we performed K-mean clustering on the dataset. To calculate the ideal number of clusters we used the elbow method. For a proper analysis, we needed at least 3 APs (two ISIs/IFRs to calculate the SD). We identified the high IFR SD cluster as bursting and the low IFR SD cluster as tonic neurons.

### Multielectrode Array

MEA data was collected with an Axion Maestro Pro multiwell MEA with a CytoView MEA24, 24-well plate and CytoView MEA48, 48-well plate (Axion Biosystems). Our iGlut culture protocol on the MEA plate is based on the culture protocol provided by Axion biosystems^45^, but we made several modifications to better fit our experimental approach. Prior to the seeding on the MEA plate, we followed the same protocol detailed above. On day 5 we coated the plates with Matrigel (4:250) for at least one hour before seeding the neurons. We dissociated the neurons from the 6 well plates following the above-described protocol and prepared a 1.2*10^7^ neuron/ml suspension. Then we placed a 10 µl droplet of the neuron suspension directly onto the electrodes, and incubated the plates at 37°C, 5% CO_2_ for 1 h. Also, we filled the space between the wells with sterile dH_2_O to prevent the cultures from drying out. After the 1 h incubation, we plated the glial cells onto the neurons (6.5-7*10^4^ cells/well) in the remaining 500 µl iGlut media (Neurobasal plus + 2% FBS). Throughout the differentiation and iGlut neuron maintenance we used the culture media compositions described above in our protocol, instead of those suggested in the Axion Cell Culture protocol. From this point, we followed our usual maintenance procedure, with half-media changes twice a week.

To analyze the MEA data, we used the AxIS Navigator (Version 3.7.2, Axion Biosystems), to detect spikes, and to make a summary of the recordings, which contains the most important information (MFR, WMFR, number of bursts and synchrony) for all wells of the MEA plate. Further, we used AxIS Metric Plotting tool (Version 2.5.0, Axion Biosystem) and Neural Metric Tool (Version 4.0.5, Axion Biosystems) to generate plots and summary reports. The WMFR is the mean firing rate, based only on those electrodes which the firing rate is greater than 5 spikes/min (active electrodes). The WMFR was only calculated for those wells which had at least one active electrode. Synchrony is defined by a synchrony index, which is a unitless value between 0 and 1^51,52^. The time window for synchrony was set as 20 ms. To calculate synchrony, a minimum of 2 spikes in a particular well was required.

### Data analysis

All subsequent data representation, analysis, and statistics were carried out in R, Python, GraphPad Prism, and Microsoft Excel software. All original code has been deposited at the GitHub repository named “High-throughput-measurements-of-neuronal-activity-in-single-human-iPSC-derived-glutamate-neurons” and is publicly available at DOI: 10.5281/zenodo.14841953 as of the date of publication. Averages are shown as mean ± SEM.

The data that supports the findings of this study are available from the corresponding author upon reasonable request.

## Author contributions

I.G.R. conceived the study, performed the Ca^2+^ imaging data acquisition and analysis, and contributed to the writing and critical revision of the manuscript. A.J.T. conceived the study, performed the MEA and patch clamp data acquisition and analysis, and contributed to the writing and critical revision of the manuscript. I.P. contributed to the initial discussions and the design of the data analysis algorithms. C.K. provided editorial feedback on the whole manuscript. Z.P.P. contributed to discussions and critical revisions of the manuscript. A.M.G. contributed to the critical revision of the manuscript. R.P.H. performed the RNAseq data acquisition and analysis, and contributed to discussions, writing, and critical revisions of the manuscript. P.A.S. conceived and supervised the study and data analyses, and contributed to the writing and critical revisions of the manuscript. All authors have approved the final manuscript.

## Acknowledgements

We want to acknowledge members of the Slesinger lab and the Goate lab for discussions throughout this work, as well as feedback from COGA investigators.

The Collaborative Study on the Genetics of Alcoholism (COGA), Principal Investigators B. Porjesz, V. Hesselbrock, A. Agrawal; Scientific Director, A. Agrawal; Translational Director, D. Dick, includes ten different centers: University of Connecticut (V. Hesselbrock); Indiana University (H.J. Edenberg, T. Foroud, Y. Liu, M.H. Plawecki); University of Iowa Carver College of Medicine (S. Kuperman, A. Anderson); SUNY Downstate Health Sciences University (B. Porjesz, J. Meyers); Washington University in St. Louis (L. Bierut, A. Agrawal, S. Hartz); University of California at San Diego (M. Schuckit); Rutgers University (D. Dick, R. Hart, J. Salvatore, J. Tischfield); The Children’s Hospital of Philadelphia, University of Pennsylvania (L. Almasy); Icahn School of Medicine at Mount Sinai (A. Goate, P. Slesinger); and Howard University (D. Scott). Other COGA collaborators include: C. Holzhauer, M. Hesselbrock (University of Connecticut); D. Lai, J. Nurnberger Jr., L. Wetherill, X., Xuei, S. O’Connor, (Indiana University); J. Kramer (University of Iowa), G. Chan (University of Iowa; University of Connecticut); C. Kamarajan, A. Pandey, D.B. Chorlian, P. Barr, S. Kinreich, G. Pandey, Z. Neale, S., C. Chatzinakos, J. Zhang, Saenz deViteri, R. Christian, A. Bingly (SUNY Downstate); G. Pathak (Icahn School of Medicine at Mount Sinai); A. Anokhin, K. Bucholz, F. Dong, A. Hatoum, E. Johnson, V. McCutcheon, J. Rice, S. Saccone (Washington University); F. Aliev, Z. Pang, S. Kuo, S. Brislin, J. Moore (Rutgers University); A. Merikangas (The Children’s Hospital of Philadelphia and University of Pennsylvania); M. Gitik, NIAAA Staff Collaborator. We continue to be inspired by our memories of Henri Begleiter and Theodore Reich, founding PI and Co-PI of COGA, and also owe a debt of gratitude to other past organizers of COGA, including Ting-Kai Li, P. Michael Conneally, Raymond Crowe, and Wendy Reich, for their critical contributions. Special thanks to the COGA collaborators who collected and classified the samples from the donors that were used in this study. This national collaborative study is supported by NIH Grant U10AA008401 from the National Institute on Alcohol Abuse and Alcoholism (NIAAA) and the National Institute on Drug Abuse (NIDA). We also gratefully acknowledge the support provided by the Training Program in Stem Cell Biology fellowship from the New York State Department of Health (NYSTEM-C32561GG) to I.G.R.

## Figure legends

**Supplementary Figure S1:**
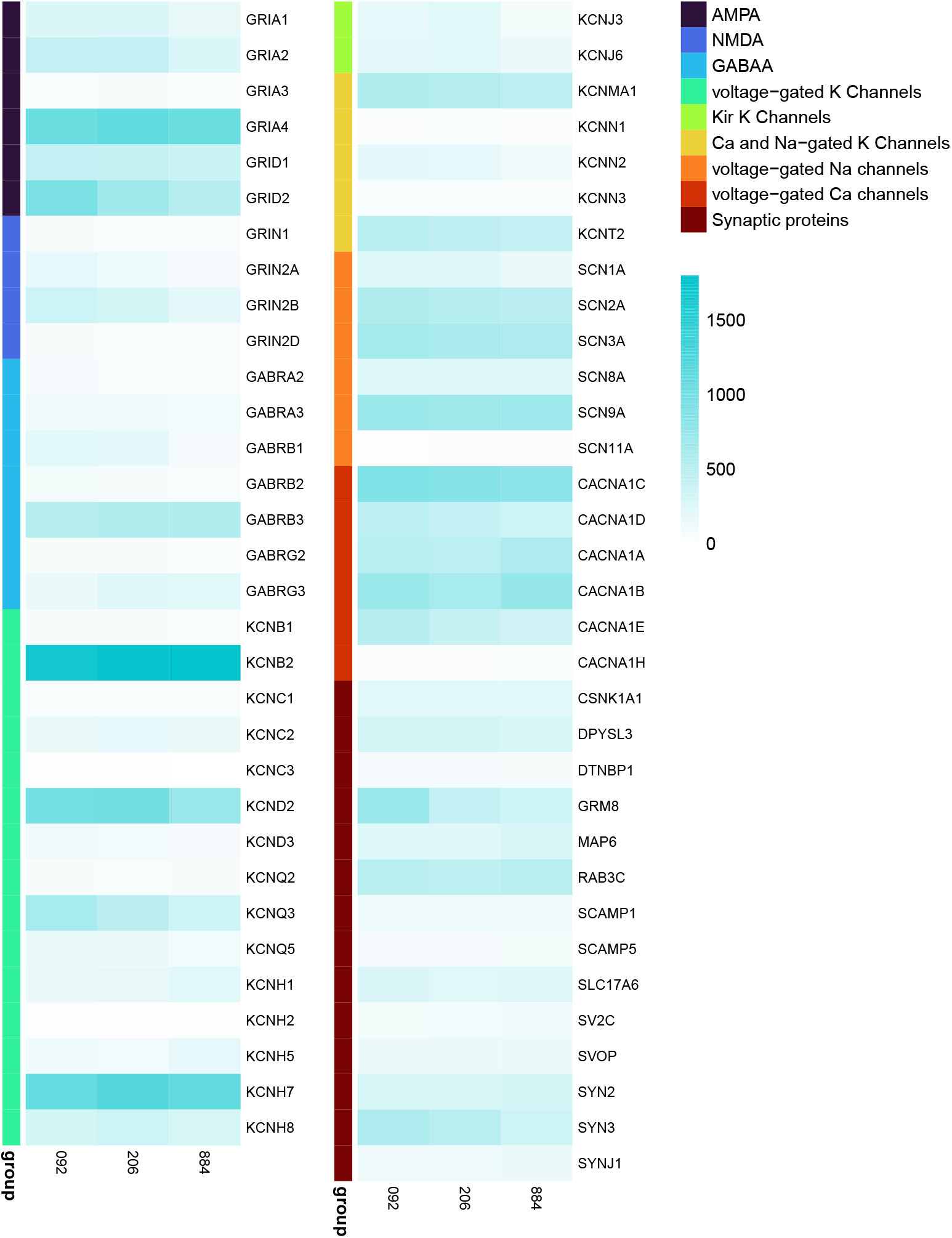
Expression levels of neurotransmitter receptor and signal transduction genes. Single-cell RNA sequencing was performed on induced neuron cell villages containing all three donor lines (10884, 9206 and 8092). mRNAs for both AMPA and NMDA receptors (GRID2, GRIA2, GluA2, GRIN2B, GRIA4) were detected in the three donor lines. Genes were grouped by function, as shown in the legend.

